# Reliable interpretability of biology-inspired deep neural networks

**DOI:** 10.1101/2023.07.17.549297

**Authors:** Wolfgang Esser-Skala, Nikolaus Fortelny

**Affiliations:** Computational Systems Biology Group, Department of Biosciences and Medical Biology, University of Salzburg, Hellbrunner Straße 34, 5020 Salzburg, Austria

**Keywords:** Interpretable deep learning, reproducible interpretability, biology-inspired deep learning, knowledge-primed neural networks

## Abstract

Deep neural networks display impressive performance but suffer from limited interpretability. Biology-inspired deep learning, where the architecture of the computational graph is based on biological knowledge, enables unique interpretability where real-world concepts are encoded in hidden nodes, which can be ranked by importance and thereby interpreted. In such models trained on single-cell transcriptomes, we previously demonstrated that node-level interpretations lack robustness upon repeated training and are influenced by biases in biological knowledge. Similar studies are missing for related models. Here, we test and extend our methodology for reliable interpretability in P-NET, a biology-inspired model trained on patient mutation data. We observe variability of interpretations and susceptibility to knowledge biases, and identify the network properties that drive interpretation biases. We further present an approach to control the robustness and biases of interpretations, which leads to more specific interpretations. In summary, our study reveals the broad importance of methods to ensure robust and bias-aware interpretability in biology-inspired deep learning.

## Introduction

Artificial neural networks and deep learning have demonstrated impressive performance in a wide range of prediction tasks from images to languages and board games^1^. Deep learning has equally impacted biomedical research, with striking successes in predicting protein structure from sequences^2^ or skin cancer from images^3^. Despite these successes, neural networks often remain “black boxes” that defy description by human-understandable terms^4^. This remains one of the critical limitations of deep learning algorithms for scientific, safety, ethical, and other reasons^5, 6^, and numerous approaches have been developed to make deep learning more interpretable^7^.

Interpretation approaches of deep learning models are primarily focused on the feature level^8–10^. In computational biology^11, 12^, a recent alternative approach utilizes prior knowledge on biological networks to influence the structure of the neural network, in so-called “visible”, “biologically-inspired”, or “knowledge-primed” neural networks^13–18^. In such biology-inspired deep learning models, hidden layers consist of nodes that correspond to biological entities, for example Gene Ontology (GO) terms^15^, Reactome pathways^17^, or signaling proteins^16^. Layers are then sparsely connected based on existing knowledge of relationships of the encoded biological entities. The architecture of biology-inspired models is thus informed by domain knowledge, which enables a unique interpretability^11, 12^: After training, a measure of importance is calculated for each node in the hidden layers (“node importance score”), which quantifies the importance of each biological entity in the network for the prediction task. For example, the P-NET model^17^ used mutation data to predict cancer state (primary versus metastatic cancer) in a network, where hidden nodes correspond to genes and Reactome pathways, thus identifying genes and pathways relevant for metastasis.

As with most computational analyses, it is critical for the above models to provide accurate interpretations^7, 8, 11, 19^, especially in view of their potential clinical relevance^17^. However, interpretation accuracy is challenging to measure due to the lack of appropriate gold standards^20^. This is in contrast to prediction accuracy, which is readily quantified using cross - validation^21^. We previously identified two key critical aspects for the reliable interpretability of knowledge-primed neural networks (KPNNs)^16^: (i) the robustness of node importance scores upon repeated training (also called “un-identifiability”^11^), and (ii) the interpretation biases induced by using biological network knowledge. To date, it is unclear whether or how robustness and bias-susceptibility affect different biology-inspired deep learning models. Indeed, broadly reviewing biology-inspired models^13–15, 17, 18, 22–41^, we found that (out of 25 models) only the BIOS model^25^ was trained in replicates and only the DTox model^23^ compared interpretations to networks trained on shuffled labels to rigorously control robustness and network biases.

Here, we analyze robustness and bias-susceptibility in the P-NET model^17^ and show that they limit interpretation accuracy. We next demonstrate how control experiments improve interpretation accuracy and thus enable more reliable interpretability. Found relevant in KPNNs and the P-NET model, our results suggest that robustness and bias-susceptibility are generalizable aspects that affect interpretation accuracy. In summary, our analyses demonstrate the impact of robustness and bias-susceptibility on interpretability and propose generalizable control algorithms, which improve reliable interpretability of biology-inspired neural networks.

## Results and Discussion

We first sought to assess whether robustness and bias-susceptibility affect interpretations beyond KPNNs and thus examined them in the P-NET model^17^, one of the most prominent examples of biology-inspired deep learning. P-NET uses a biology-inspired architecture of 9,229 genes and 3,073 curated biological pathways over 6 layers, which were trained on genomic data of 1,013 prostate cancer patients to predict whether patients developed metastasis. After training, P-NET uses DeepLIFT to obtain the importance scores for hidden nodes, which are ultimately used as interpretations. To assess the reliability of P-NET interpretations, we downloaded the algorithm and reproduced the original network and importance scores using the provided random seeds. Next, we assessed robustness by repeatedly training networks with varying initial weights and assessed bias-susceptibility by training on artificial control data. Finally, we corrected importance scores to yield bias-free and more specific interpretations.

### Repeated network training measures the robustness of interpretations

Robustness of node importance scores refers to the observation that repeated training of a given network on the same data can result in different interpretations in the obtained “replicate networks”, even if they achieve similar prediction accuracy. This lack of consistency is due to the random initiation of weights, which enables a network to “choose” among multiple informative input features. This in turn propagates to differences in trained weights and, ultimately, importance scores. To estimate the thus caused uncertainty of interpretations, multiple networks with the same network structure, input, and output need to be trained, resulting in a distribution of importance scores^16, 42^.

To assess the robustness of P-NET, we retrained the model 50 times with different initial weights (**Figure 1a**) by varying the random seed prior to training (see methods for details) and thereby obtained importance scores as in the original network (**Figure S1**). We then compared our replicate networks to the original network, which was originally published based on one specific random seed. From the replicate networks, we obtained a distribution of importance values for every node. Notably, importance scores from replicate networks are equally plausible as those from the original network, because (i) the random seed is neither a meaningful nor tunable (hyper)parameter, and (ii) all replicate networks had comparable predictive power to the original network (**Figure 1b**). For many nodes, the importance scores from replicate networks were similar to the importance scores from the original network. However, in a significant number of cases, the original scores diverged from the scores from replicate networks (**Figure 1c, d**). For example, in layers 5 and 6 the most important node of the original network was ranked only second in the replicate networks.

**Figure 1.**
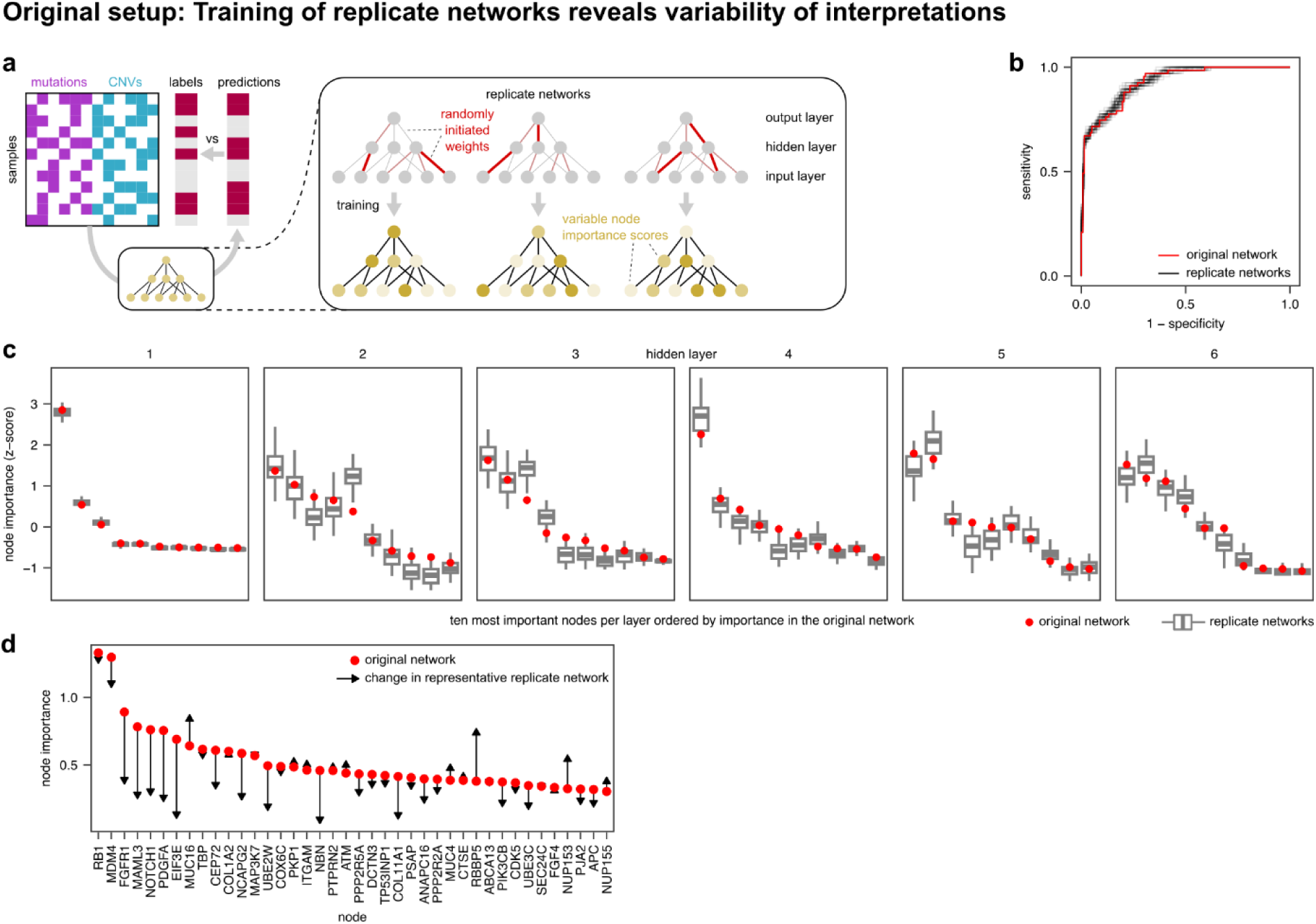
Robustness of interpretations of the P-NET model. (a) Outline of the experimental approach. (b) ROC curves for the original setup and replicate networks. (c) Node importance in replicate networks (n = 50 replicate networks; trained with different random seeds for weight initiation) compared to original importance (trained with seeds from the published model). (d) Change of node importance scores in one representative replicate network (black lines) compared to the scores in the original network (red dots).

Importantly, these observations are not limited to P-NET and KPNNs. To demonstrate the broad relevance of assessing robustness, we analyzed interpretations in the DTox model^23^ (**Figure S2**). Training replicates of DTox, we found a wide spread of correlations between replicate networks, ranging from highly positive (maximum R = 0.98) to highly negative (minimum R = -0.98) correlations, which shows a limited robustness.

Taken together, our analyses of the P-NET and DTox models demonstrate that interpretations are affected by the choice of initial weights, which are randomly and thus arbitrarily selected. In line with previous research^16, 42^, our results show that repeated network training is required to assess the robustness of interpretations in different biology-inspired models and that averaged interpretations of replicate networks provide more reliable interpretations than individual networks^11^.

### Control inputs reveal interpretation biases under high prediction accuracy

Biological networks are highly biased with very few highly connected nodes (hubs) and many nodes with fewer connections^43^, which can influence importance scores of biology-inspired neural networks. In our previous work on KPNNs, we found that hubs tend to get larger importance scores than less connected nodes, irrespective of the input data, since hubs have access to more input features and thus more information^16^.

To assess whether network biases affect importance scores in P-NET, we used “deterministic” control inputs, an approach we previously developed for KPNNs^16^. These control inputs are artificially designed such that every input feature is perfectly correlated with the target labels and can thus perfectly predict the labels (**Figure 2a**). Since all control features are equally informative, the differences in importance scores between nodes (after training) are solely driven by the network structure and not by the data. Training P-NET on our deterministic inputs, we observed clear differences of importance scores between nodes (**Figure 2b**), demonstrating that some nodes receive high importance scores purely based on network biases. Importantly, these scores were reproducible across 50 replicate networks trained on deterministic inputs, demonstrating that the observed differences in importance scores reveal biases inherent to the prediction task, and not random variability that may arise from random weight initialization. Deterministic control inputs thus enabled us to robustly assess the influence of network biases on node importance scores in the extreme case of an “easy” prediction task with a very high prediction performance (i.e., an AUC of 1; **Figure 2c**).

**Figure 2.**
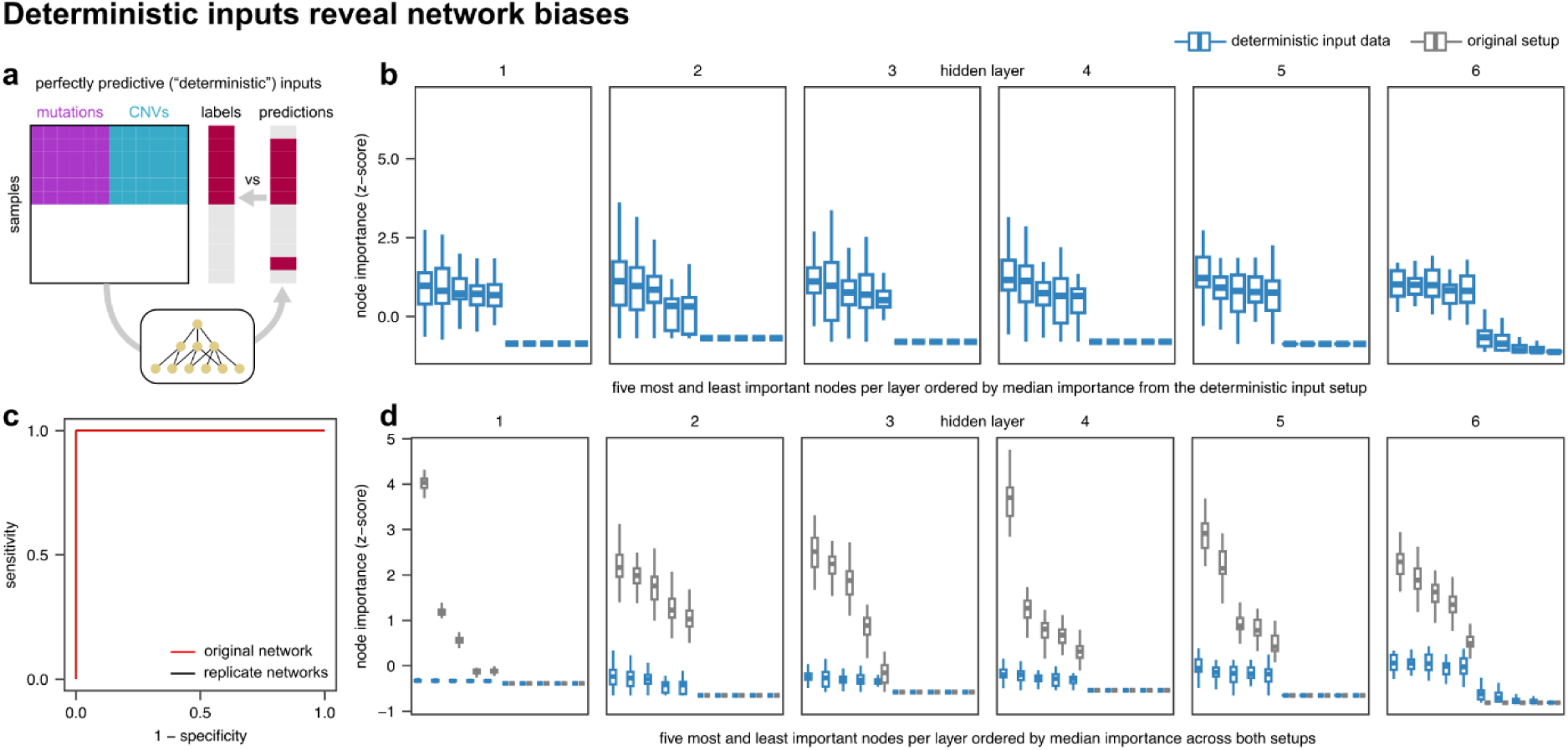
Network biases in the P-NET model based on deterministic input data. (a) Experimental approach. (b) Node importance derived from deterministic inputs (n = 51 replicate networks), where every input feature is perfectly correlated with the target labels. (c) ROC curves for the original setup (from Figure 1) and replicate networks. (d) Comparison of importance scores for deterministic (n = 51 replicate networks) and original input data (n = 51 replicate networks).

We next assessed whether importance scores obtained from training on original input data differed from those obtained from training on deterministic control inputs (**Figure 2d**). Indeed, importance scores from real data greatly exceeded those from control inputs for nodes with high importance scores. The resulting large “differential” node scores (obtained from comparing importance scores from original data to deterministic inputs for each node) suggest that the top nodes are important beyond what is expected from network biases alone and are thus reliable interpretations. However, beyond the top nodes, we found many nodes with control importance scores that equaled or exceeded real importance scores, indicating an inflated importance measure that is mainly driven by network biases. In summary, our deterministic control inputs revealed the effect of network biases on importance scores and enabled a differential analysis that corrects for the observed biases.

### Label shuffling reveals biases under low prediction accuracy

While our deterministic control inputs rely on perfectly predictive inputs, we next sought to examine biases under the opposite case, i.e. networks with limited predictive power. To this end, we randomly shuffled output labels, an approach commonly used to assess biases in prediction performance such as class imbalance (**Figure 3a**). We shuffled target labels before training, such that the prediction algorithm should only be able to learn spurious relationships in the training data (i.e., an AUC close to 0.5 on the test data). With regards to interpretability, shuffled labels complement deterministic control inputs: While interpretations from deterministic inputs (**Figure 2**) reveal network biases under high prediction accuracy, interpretations from shuffled labels reveal biases under low prediction accuracy. After training the P-NET model on shuffled labels, we again observed a distribution of importance measures across nodes (**Figure 3b**) and an expectedly low prediction performance on test data (AUC around 0.5; **Figure 3c**). Similar to the node importance scores obtained from deterministic inputs (**Figure 2**), importance scores from shuffled labels were also reproducible across replicate networks (**Figure 3c**), demonstrating that node importance scores reflect network biases and are not random. In contrast to deterministic inputs, we found that importance values from shuffled labels were on a similar scale to those from original labels (**Figure 3d**). Therefore, for some of the top nodes (with highest original importance scores), the high importance scores from shuffled labels suggested a large influence of network biases.

**Figure 3.**
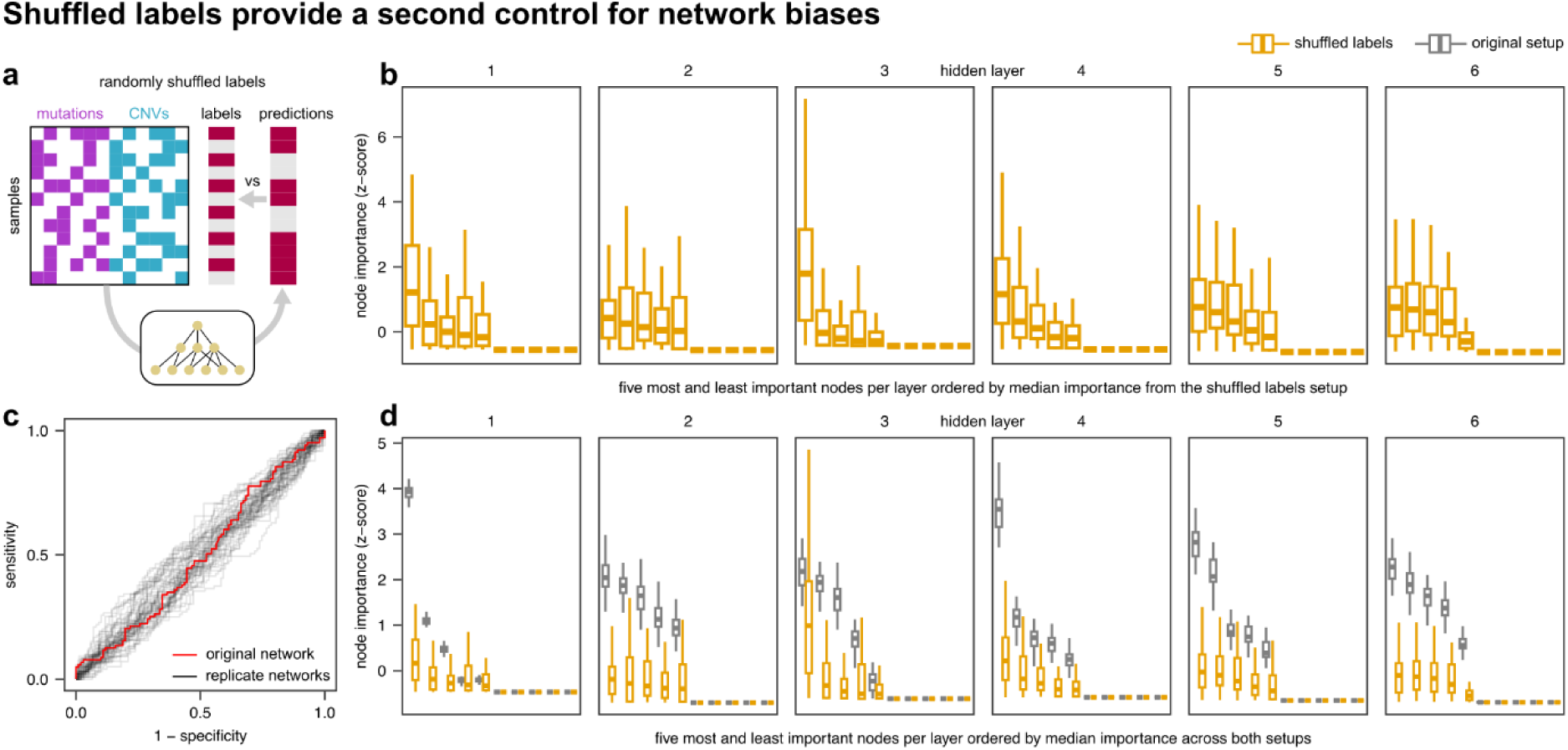
Network biases in the P-NET model based on shuffled target labels. (a) Experimental approach. (b) Node importance derived from shuffled labels (n = 51 replicate networks). (c) ROC curves for the original setup (from Figure 1) and replicate networks. (d) Comparison of importance scores for shuffled (n = 51 replicate networks) and original labels (n = 51 replicate networks).

Taken together, shuffled labels and deterministic control inputs provide two approaches to assess network biases from both angles: low and high prediction accuracy. To better understand similarities and differences of these approaches, we compared importance scores across the three experimental setups (original setup, deterministic inputs, or shuffled labels; **Figure 4**), which confirmed that node importance scores were reproducible across replicate networks within each setup. Interestingly, scores calculated using shuffled labels were more similar to scores obtained in the original setup (Pearson’s R = 0.48) than scores derived from deterministic inputs (Pearson’s R = 0.25). These observations were consistent across all layers (**Figure 4**) and within each layer (**Figure S3**).

**Figure 4.**
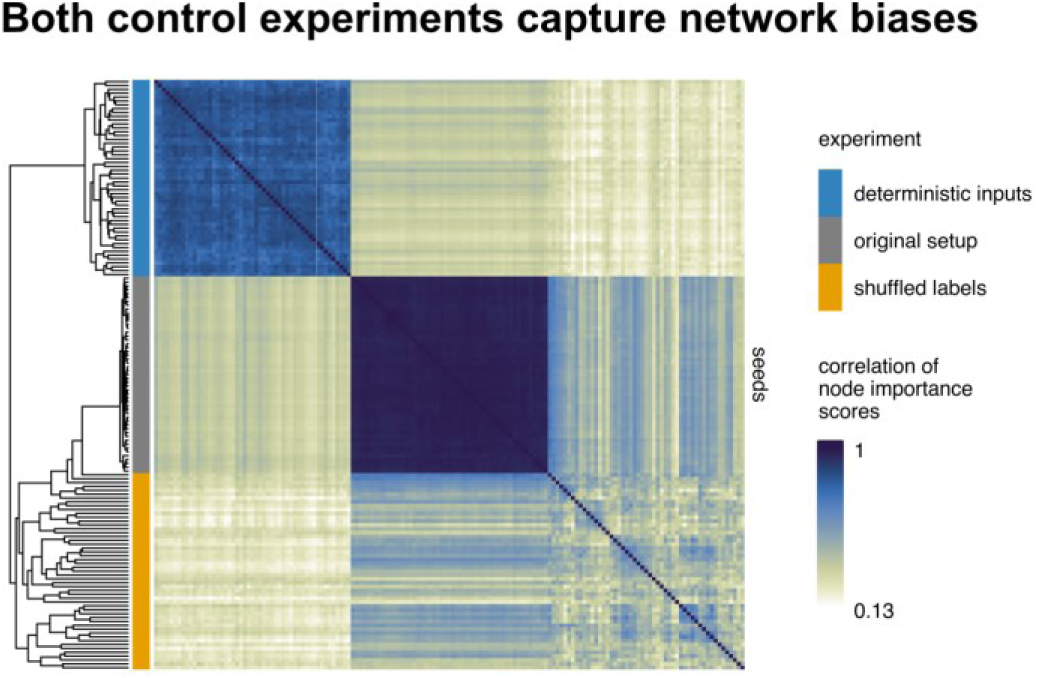
Correlation (Pearson’s R) of node importance scores from all layers across experimental approaches. Each of the 103 rows and columns (51 seeds x 3 setups) represents a different network.

### Control experiments capture network biases

As both control approaches yielded reproducible but different importance scores, we next sought to assess which network properties may be captured by each approach. We correlated importance scores to measures of node centrality, which provide network-based and node-level measures of network biases (**Figure 5, Figure S4**). Indeed, we found that importance scores were highly correlated with network reachability and betweenness in the original P-NET setup and in the setup using shuffled labels. Node reachability measures the number of nodes from a given node is “reachable” through paths of any length^44^. In a neural network, reachability can be interpreted as the amount of information available to a given node^16^. Betweenness measures the number of shortest paths between all pairs of nodes that path through a given node. Both measures rely on paths through the network, thus measuring “global” node properties. Interestingly, both measures were also highly correlated (**Figure S5**), which is not a general observation^44^ but likely due to the particular architecture of feed-forward neural networks^16^.

**Figure 5.**
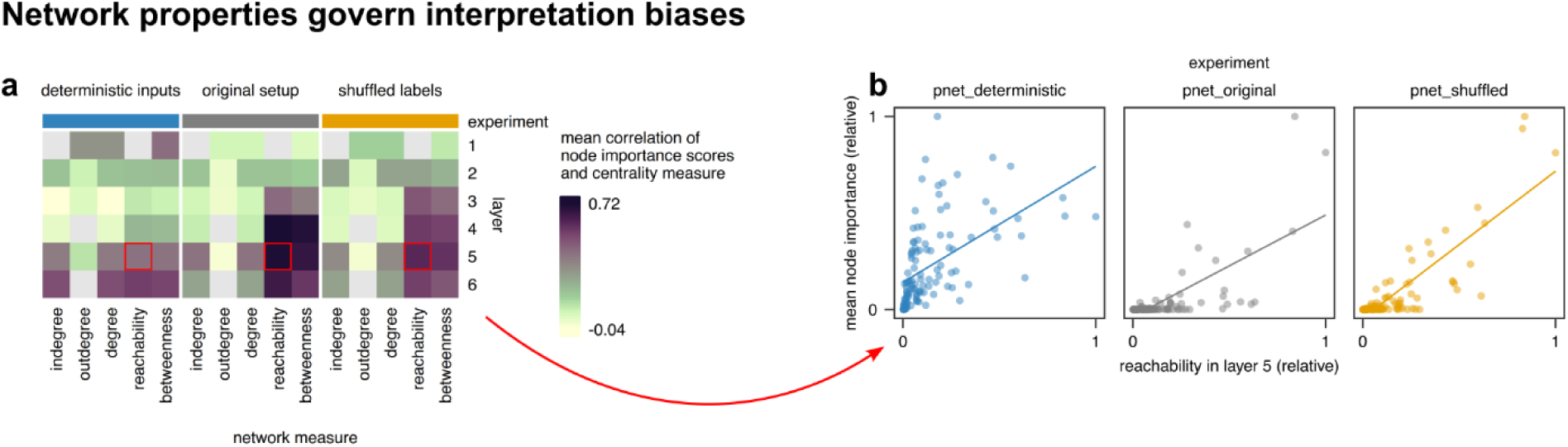
(a) Correlation (Pearson’s R) of node importance scores and network centrality measures. Gray denotes missing values. Indegree and outdegree correspond to the number of nodes to which a given node is connected in the previous and following layer, respectively. Reachability denotes the number of nodes from which a node is reachable. The betweenness is related to the number of shortest paths between all pairs of nodes that pass through a node. Notably, some layers contain nodes with a single value for the centrality measure. For instance, all nodes in layer 1 have indegree 3 (three predecessors with mutation, copy number amplification, and copy number deletion data) and thus a reachability of 4 (since the node is reachable from its predecessors and from itself). (b) Exemplary scatterplots for the cells highlighted in (a); each point corresponds to a hidden node (n = 113 in each panel).

In contrast to the above global measures, importance scores were less correlated with node degree, which measures the direct neighbors of each node and is thus a local measure. Of note, P-NET implements a normalization of importance scores based on node degree in order to attenuate importance measures of highly connected nodes. Our results, which were obtained after this normalization, demonstrate that a normalization based on local network measures does not correct for network biases of interpretations.

### Correction approach yields more reliable and specific interpretations

To obtain interpretations that account for robustness and network bias, we calculated differential importance scores by comparing importance scores from the original setup to the control of shuffled labels (**Figure 6**), thus comparing the original scores to a background. A positive differential score indicates that a node is more important than expected from network biases alone. A score close to zero or a negative score suggests that importance is either driven by network biases or less than expected by these biases, respectively. We focused our analyses on the first (gene) layer, which was also the focus of interpretations in P-NET^17^: First, AR has a highly positive differential score, suggesting that this gene is much more relevant than expected from network biases. This highlights the important and highly specific role of androgen signaling in prostate cancer, which is supported by numerous publications^45–47^. Second, TP53 received a negative differential score, indicating that the importance is driven by network biases. Indeed, TP53 is a highly multifunctional gene and known to bias computational analyses^48^. Third, MDM4 had a positive differential score, showing that this gene is specific to the process studied, which is supported by the experiments performed in the P-NET publication^17^. Differential importance scores thus suggest a much more specific and critical role of androgen signaling (AR) in pancreatic cancer compared to TP53, which is in line with existing literature^45–47^.

**Figure 6.**
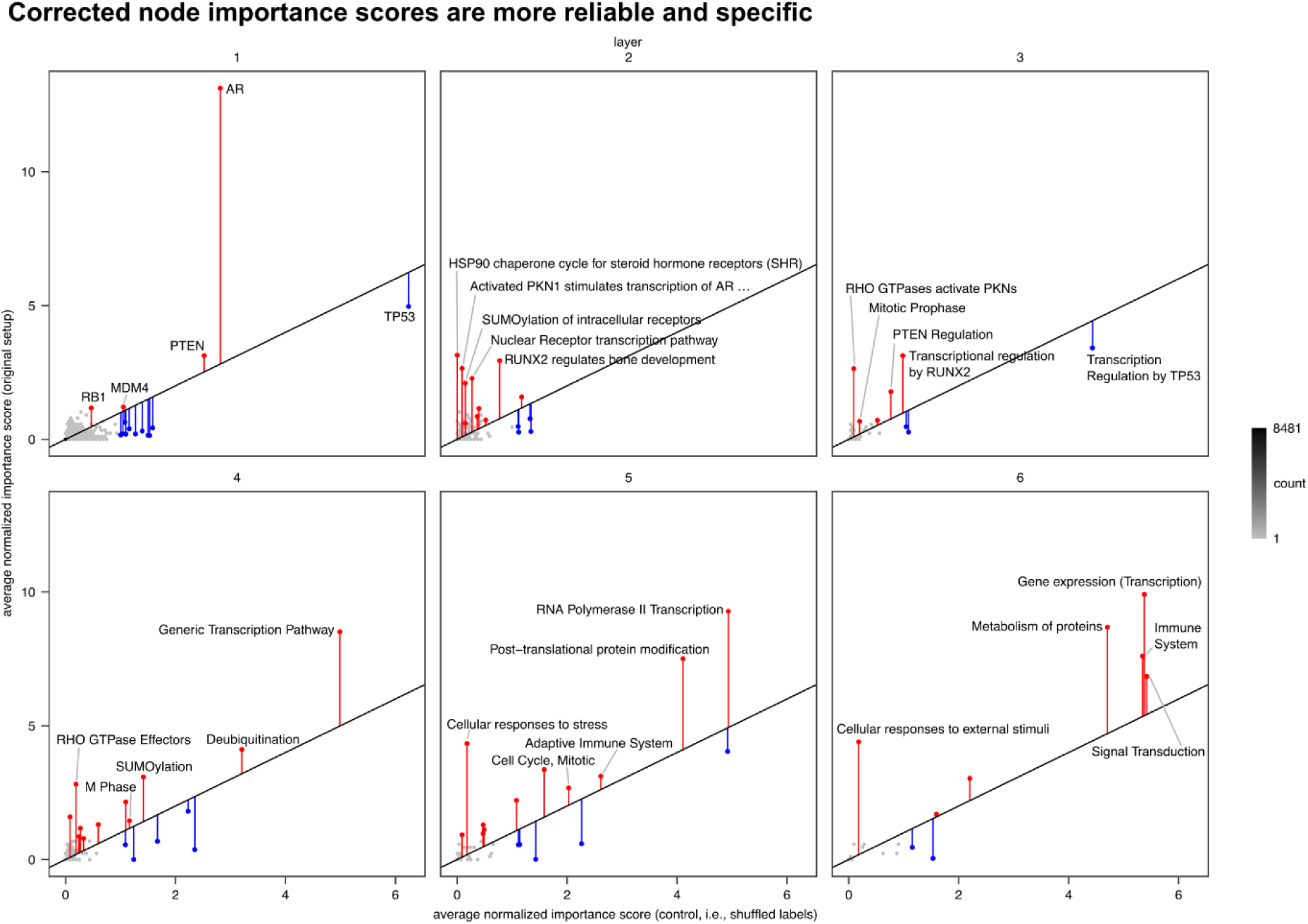
Correction approach in the P-NET model. Raw importance scores (original setup) shown on the y-axis are compared to control importance scores (shuffled labels) on the x-axis. Vertical bars are shown for nodes with high scores (i.e., scores greater than one in either approach) and show differential importance scores. In each layer, the five nodes with the highest importance score in the original setup using the original seed are labeled.

To further validate the increased specificity obtained by correcting node interpretations, we trained the P-NET model on the MSK-IMPACT 2017 dataset^49^ (see methods for details), a dataset that contains genomic data similar to the data used in original P-NET model but from different cancers (**Figure S6**). We trained replicate P-NET networks on genomic data from four cancers (non-small cell lung, breast, colorectal, and prostate) to predict metastasis and then compared node importance scores between cancers. Interestingly, when comparing raw importance scores, we found a strong correlation (from R = 0.55 to R = 0.83) between cancers, demonstrating that interpretations were not specific to the cancer and input data but likely driven by other factors such as network biases. In contrast, after correcting importance scores by calculating differential scores, correlations between different cancers significantly decreased (from R = -0.02 to R = 0.47). This demonstrates the ability of our correction procedure to remove biases from interpretation scores, resulting in corrected scores that are more specific to the dataset (or here cancer) studied. Taken together, our results from the original P-NET dataset and from MSK-IMPACT 2017 show that our correction approach corrects interpretations, removing biases and revealing effects that are specific to the biological process studied.

## Conclusion

Biology-inspired neural networks enable a unique type of interpretability by incorporating domain knowledge of biological relationships into the architecture of deep learning models^11, 12^. However, critical steps must be taken to ensure reliability of interpretations. Our results demonstrate that robustness and network-biases, which we previously studied in KPNNs^16^, broadly affect biology-inspired models. We further demonstrate that control methods can improve the accuracy of interpretations but are lacking in the field of biology-inspired deep learning.

Controlling robustness, we repeatedly trained identical networks on identical data, which reveals that interpretations vary based on the choice of randomly initiated weights. Similar observations have been made for autoencoders^42^ and can likely be expected for any algorithms relying on random initial parameters. Robustness towards small changes in the model or data has been widely recognized is key for machine learning^19^. For example, in graph neural networks, “stability” is quantified upon small perturbations of nodes or edges^50^. Here, we demonstrated that dedicated analyses are required to ensure robustness for biology-inspired neural networks.

Controlling for network biases, we used deterministic inputs and shuffled labels. These analyses revealed that, without controls, certain nodes receive higher importance scores based on their position in the network but independently of input data and output labels. Network biases specifically affect biology-inspired neural networks, where interpretations are focused on hidden nodes, and have not been described for other interpretation algorithms focused on input features. However, similar biases have been shown to bias gene function predictions in biological networks^48^ and thus likely represent a general aspect of working with biological networks.

Taken together, we have confirmed challenges for interpretability in the P-NET model^17^ and have described and characterized control experiments. We expect that these findings will also be relevant for other biology-inspired neural networks or related machine-learning interpretation methods.

## Methods

### Preparation of P-NET for manual selection of random seeds

P-NET uses two hard-coded random seeds to ensure reproducible network training. In order to facilitate manual selection of these seeds, the P-NET code was downloaded from GitHub (https://github.com/marakeby/pnet_prostate_paper, commit 2b16264 dated 2021-11-15), and the files pipeline/one_split.py and train/run_me.py were modified. These patches allowed to specify two seeds as command line arguments when running the latter script. The Python environment (comprising Python v2.7.15 and TensorFlow v1.12.0) was set up via conda (v4.10.3) through the environment.yml file provided in the P-NET repository. In order to facilitate reproducible analysis across platforms, a Docker container with P-NET installed is available from our GitHub Container registry (ghcr.io/csbg/pnet-container).

### Training of replicate networks in the P-NET model

In each of the three experimental setups described below, P-NET was trained repeatedly, once with the original random seeds (234 and 20080808) and 50 times with seeds ranging from 0 to 49. As the value of the random seed influences the choice of initial weights, this enables us to assess robustness of interpretations. The number chosen (50 replicate networks) enabled us to estimate a distribution of importance scores (for the robustness analysis in **Figure 1**) and was sufficient to show clear differences in importance scores of interpretations from the original setup and the control experiments (**Figures 2–3**). Training was conducted on an HP EliteBook 850 G7 containing an Intel Core i5-10210U CPU (4 cores, 1.6 GHz). In all setups, the training, validation, and test set comprised 80%, 10%, and 10% of the data, respectively. (a) The original setup utilized the P-NET input data files available for download (https://drive.google.com/uc?id=17nssbdUylkyQY1ebtxsIw5UzTAd0zxWb). (b) Deterministic control inputs were obtained by modifying two P-NET input data files: _database/prostate/processed/P1000_final_analysis_set_cross_important_only.csv, a binary matrix describing the presence (1) or absence (0) of at least one mutation in 14378 genes of 1011 samples; and _database/prostate/processed/P1000_data_CNA_paper.csv, a decimal matrix describing copy number alterations in 13802 genes of 1013 samples. (In the latter matrix, P-NET interpreted values above 1.5 as presence of copy number amplification and values below –1.5 as presence of copy number deletion). These matrices were modified such that in all samples labeled as metastatic, each gene was characterized by mutation (value 1) and copy number amplification (value 2), while in the samples labeled as normal, each gene received values denoting wild type (value 0) and normal copy number (value 0). (c) Shuffled labels were obtained by randomly labeling all 1013 samples listed in _database/prostate/processed/response_paper.csv as either metastatic (value 1) or normal (value 0) while ensuring equal class frequency.

### Training of replicate P-NET networks on MSK-IMPACT 2017 data

P-NET was trained repeatedly (n = 51, as described above) on the MSK-IMPACT 2017 dataset^49^ in two experimental setups: (a) The original setup utilized labels (primary or metastatic), mutations (detected in 414 genes), and copy number variations (detected in 410 genes) from the dataset. (b) Shuffled labels, with the same input data but shuffled output labels as described above for the original P-NET dataset. Both experimental setups were applied to the four most frequent cancer types in the dataset: non-small cell lung cancer (1668 samples), breast cancer (1337 samples), colorectal cancer (1007 samples), and prostate cancer (717 samples).

### Training of replicate networks in the DTox model

DTox^23^ uses one hard-coded random seed to ensure reproducible network training. In order to facilitate manual selection of this seed, the DTox code was downloaded from GitHub (https://github.com/EpistasisLab/DTox, commit 10c909b dated 2022-10-13), and the files code/dtox.py and code/dtox_learning.py were modified. These patches allowed to specify a seed when running DTox and to export predicted labels. The Python environment (comprising Python v3.7.3 and torch v1.10.1) was set up via conda (v23.3.1) through the environment.yml file provided in the DTox repository. A Docker container with DTox installed is available from our GitHub Container registry (ghcr.io/csbg/dtox-container).

DTox was trained repeatedly, once with the original random seed (0) and 50 times with seeds ranging from 1 to 50. All other settings were kept consistent with the original DTox publication: To train and test each network, we used the dataset available in the DTox GitHub repository (in the folder data/example). Both the training and test set comprised 500 compounds each (samples), characterized by a 166-bit MACCS fingerprint, from which binding probabilities for 361 target proteins were calculated as a preprocessing step. Node importance scores were calculated for those 97 compounds that showed activity in a mitochondria toxicity screen.

### Data analysis

All analyses were conducted in R (v4.3.1)^51^. ROC curves were calculated by the ‘roc_curve’ function in yardstick (v1.2.0)^52^. Node importance scores from all layers across experimental approaches were compared by calculating Pearson correlation coefficients *r_ij_* (R function ‘cor’) and using these coefficients as distances 1 – *r_ij_* for hierarchical clustering (R function ‘hclust(method = “complete”)’). Network centrality measures were calculated by igraph (v1.4.3)^53^. Figures were plotted with ggplot2 (v3.4.2)^54^ and ComplexHeatmap (v2.16.0)^55^. To calculate differential importance scores, we normalized scores using the function ‘normalizeQuantiles’ from limma (v3.56.2), and subtracted control importance scores from those from the original setup. Shuffled labels were used as the control and scores from replicate networks were averaged for each node.

### Code availability

Code for setting up and running P-NET, analyzing its output, and generating all figures is available from GitHub (https://github.com/csbg/pnet_robustness).

### Data availability

Data used for generating all figures is available from https://doi.org/10.5281/zenodo.7760561.

## CRediT author statement

WES: Methodology, Software, Investigation, Resources, Data Curation, Visualization, Writing - Original Draft, Writing - Review & Editing; NF: Conceptualization, Methodology, Investigation, Writing - Original Draft, Writing - Review & Editing, Supervision, Project administration.

---

**Figure S1.**
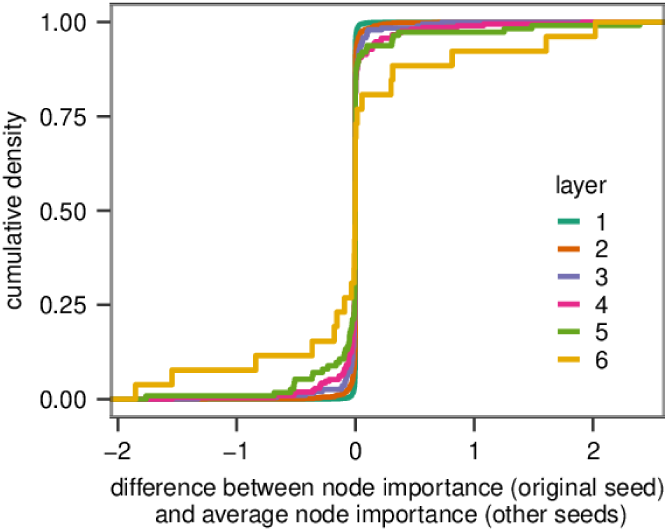
Distribution of the differences between node importance scores obtained using the original seed and the average node importance seed across the 50 replicate networks (n = 11,147 differences).

**Figure S2.**
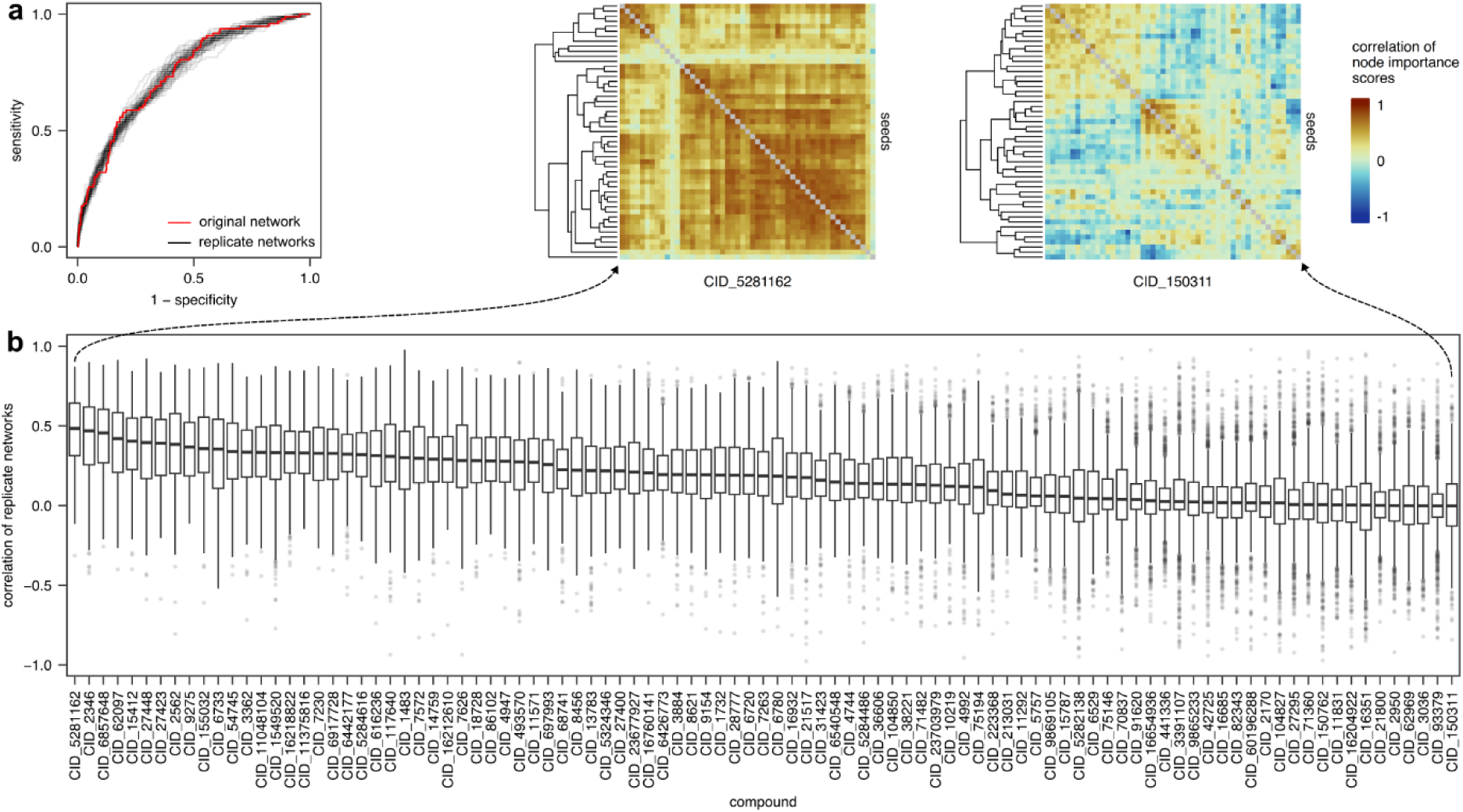
Analysis of robustness in the DTox model^23^. DTox predicts toxicity (output labels) of compounds (observations) from their structural properties (input features) in a network constructed from Reactome pathways, providing interpretations at the level of proteins and pathways. Importantly, interpretations of DTox are local interpretations in that one set of importance scores is obtained for every compound (observation) and trained replicate network. (a) ROC curves of replicate DTox networks (n = 51 replicate networks), showing that prediction performance is comparable between replicates. (b) Pairwise correlations (Pearson’s R) between replicate networks across importance scores obtained from DTox, showing a wide range of correlations and thus a limited robustness for a large number of compounds (n = 1,275 correlation values per compound) For the compounds with the highest and lowest median correlation, heatmaps depict correlations (Pearson’s R) of the replicate networks.

**Figure S3.**
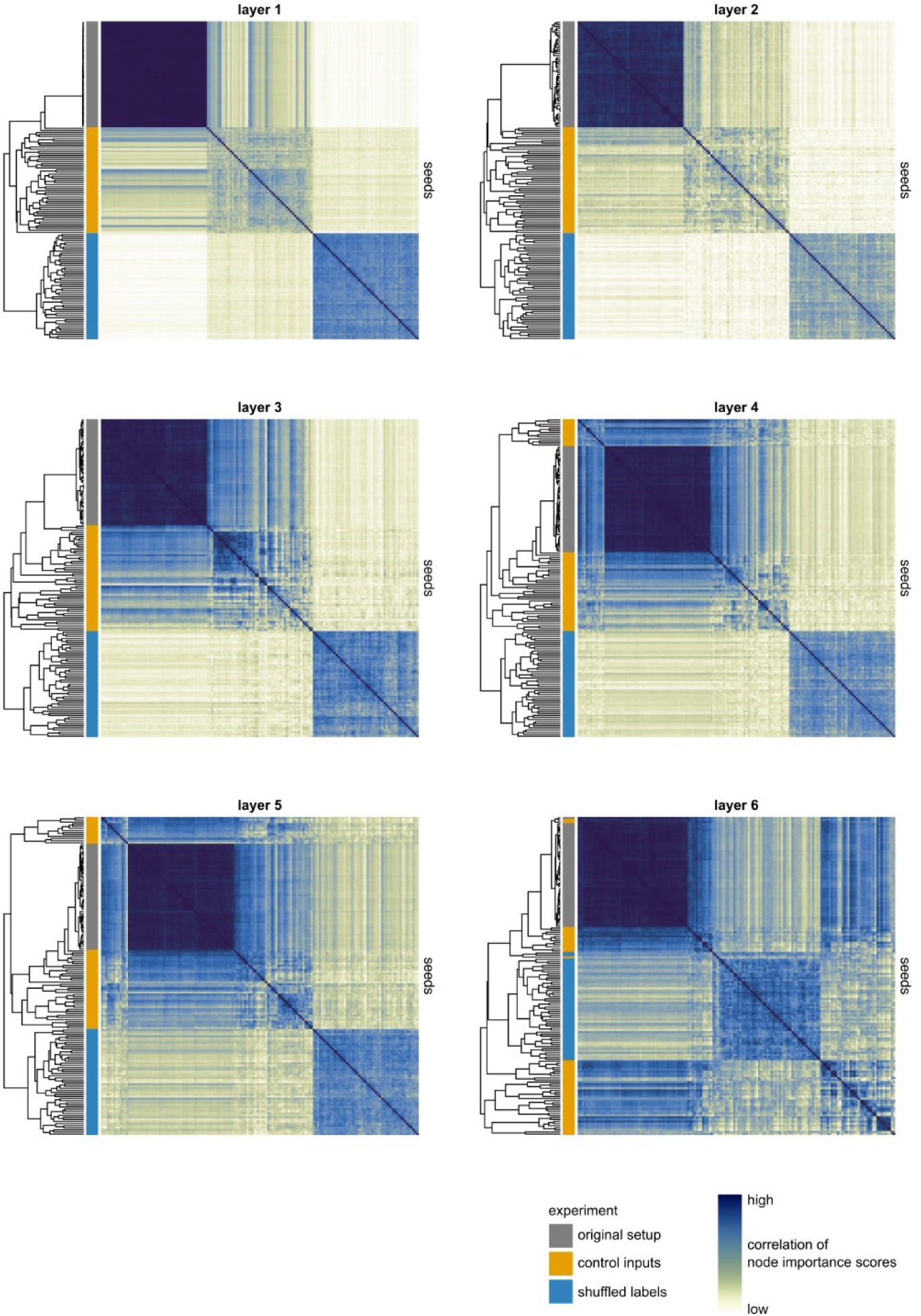
Layer-wise correlation (Pearson’s R) of node importance scores across experimental approaches. Each of the 103 rows and columns (51 seeds x 3 setups) represents a different network.

**Figure S4.**
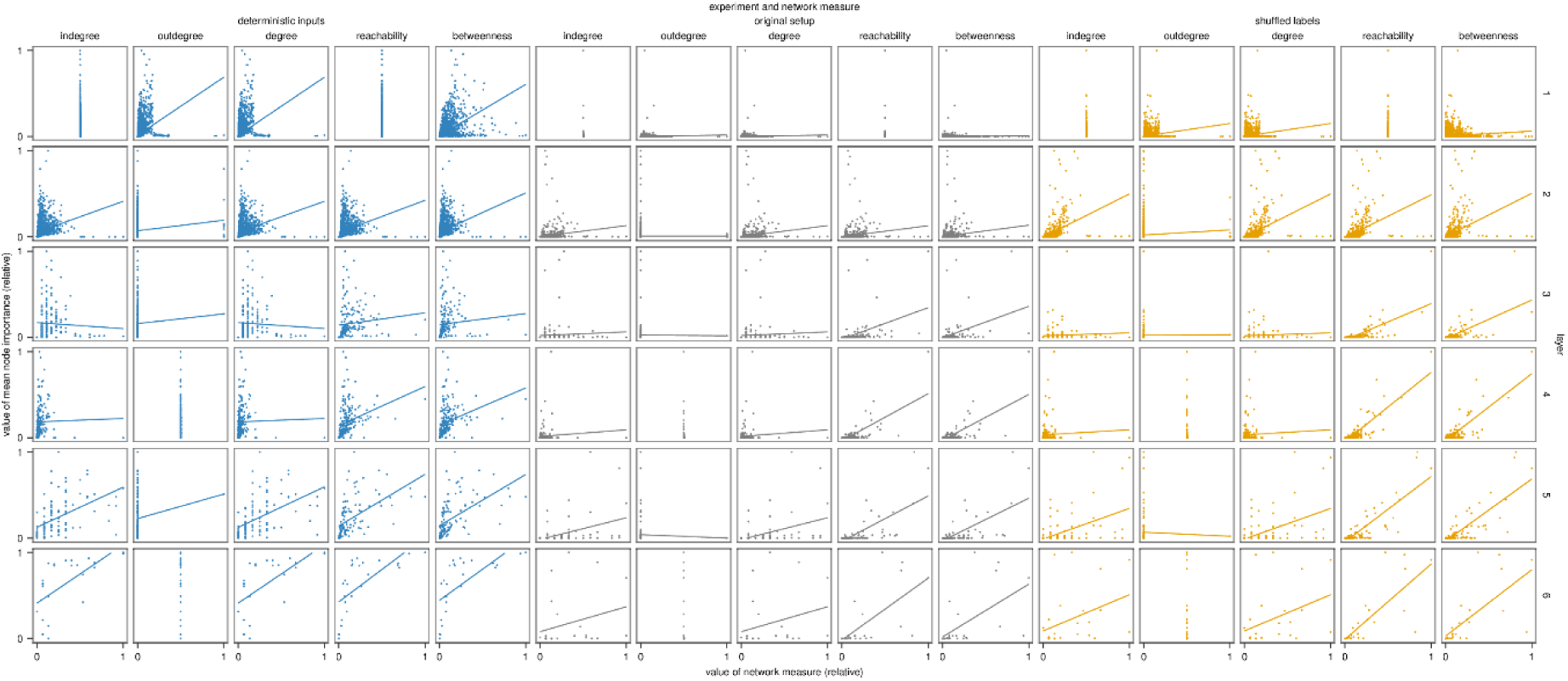
Scatterplots for all cells of the heatmap in Figure 5a. Each point corresponds to a hidden node (n = 9229, 1387, 193, 210, 113, and 26 in each panel of layers 1 to 6, respectively).

**Figure S5.**
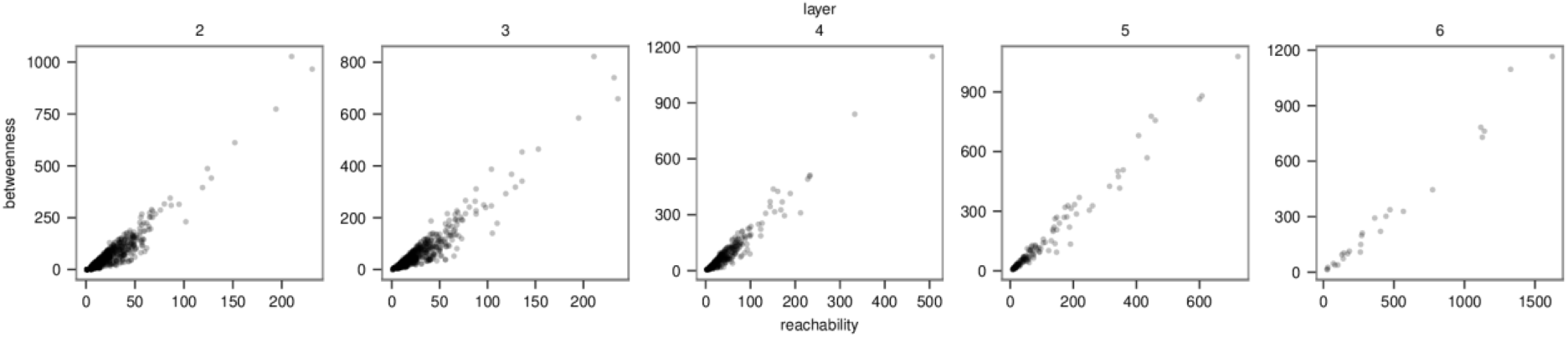
Scatterplot of betweenness and reachability in each P-NET layer. The subplot for layer 1 (gene layer) is omitted, since all nodes in this layer share a reachability of 1.

**Figure S6.**
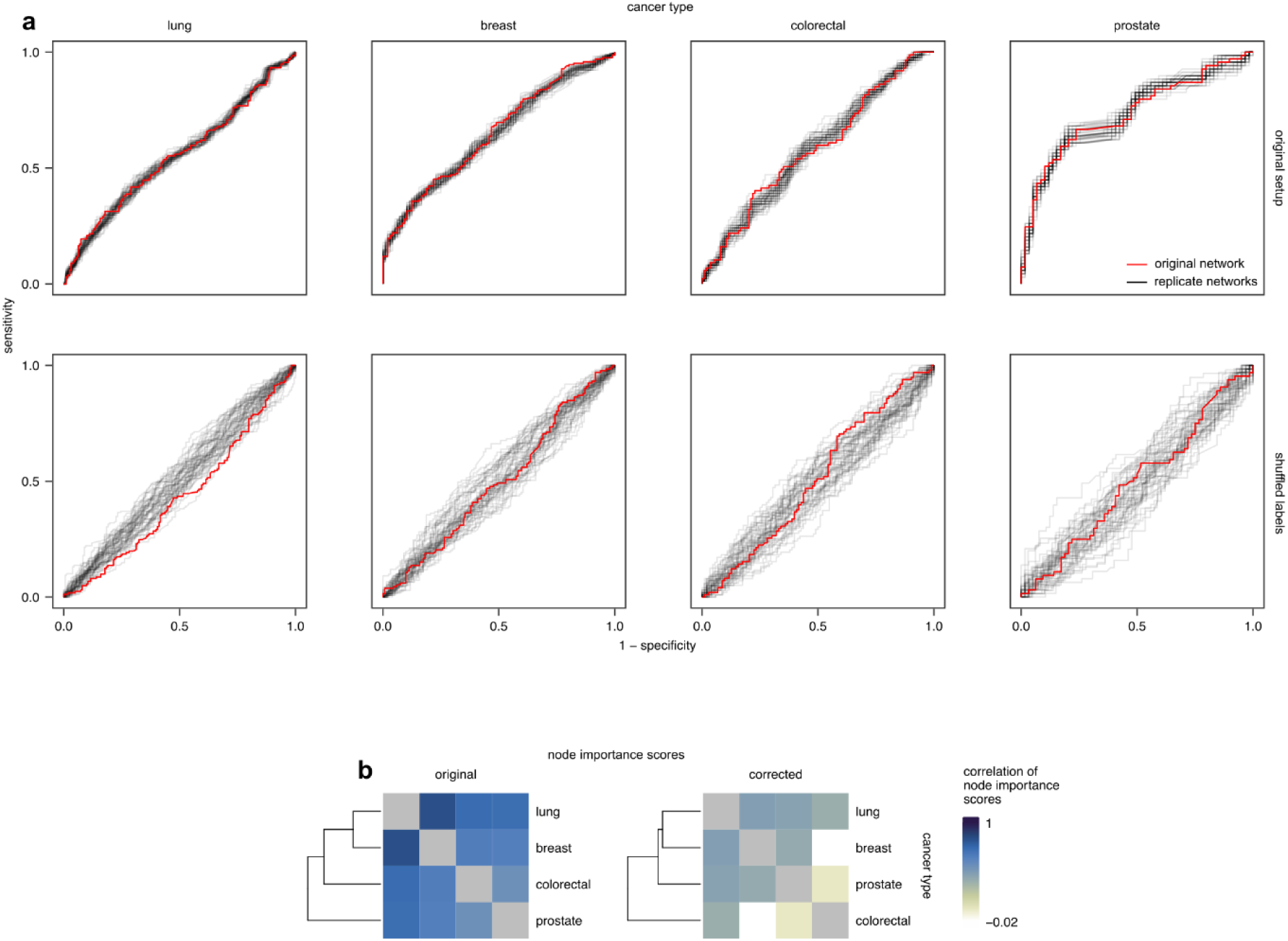
Correction approach on MSK-IMPACT data. (a) ROC curves of P-NET trained on subsets (i.e., different cancer types) of the MSK-IMPACT 2017 dataset (n = 51 replicate networks per experiment). ROC curves are shown for the original setup (top) and shuffled labels (bottom). (b) Correlation (Pearson’s R) heatmap showing correlations of average node importance scores from the original setup (left) and after the correction (right) using differential importance scores.

